# Cross cell-type systems genetics reveals the influence of eQTL at multiple points in the developmental trajectory of mouse neural progenitor cells

**DOI:** 10.1101/2025.01.24.634514

**Authors:** Selcan Aydin, Daniel A. Skelly, Hannah Dewey, J. Matthew Mahoney, Ted Choi, Laura G. Reinholdt, Christopher L. Baker, Steven C. Munger

## Abstract

Genetic variation leads to phenotypic variability in pluripotent stem cells that presents challenges for regenerative medicine. Although recent studies have investigated the impact of genetic variation on pluripotency maintenance and differentiation capacity, less is known about how genetic variants affecting the pluripotent state influence gene regulation in later stages of development. Here, we characterized expression of more than 12,000 genes in 127 donor-matched Diversity Outbred (DO) mouse embryonic stem cell (mESC) and neural progenitor cell (mNPC) lines. Quantitative trait locus (QTL) mapping identified 2,947 expression QTL (eQTL) unique to DO mNPCs and 1,113 eQTL observed in both mNPCs and mESCs with highly concordant allele effects. We mapped three eQTL hotspots on Chromosomes (Chrs) 1, 10, and 11 that were unique to mNPCs. Target genes of the Chr 1 hotspot were overrepresented for those involved in mRNA processing, DNA repair, chromatin organization, protein degradation, and cell cycle. Mediation analysis of the Chr 1 hotspot identified *Rnf152* as the best candidate mediator expressed in mNPCs, while cross-cell type mediation using mESC gene expression along with partial correlation analysis strongly implicated genetic variant(s) affecting *Pign* expression in the mESC state as regulating the mNPC Chr 1 eQTL hotspot. Together these findings highlight that many local eQTL confer similar effects on gene expression in multiple cell states; distant eQTL in DO mNPCs are numerous and largely unique to that cell state, with many co-localizing to mNPC-specific hotspots; and mediation analysis across cell types suggests that expression of *Pign* early in development (mESCs) shapes the transcriptome of the more specialized mNPC state.

## Introduction

Pluripotent stem cells (PSCs) have the capacity to self-renew indefinitely in culture and differentiate into any cell type. These properties make them a valuable resource for the study of gene regulation in various cellular contexts, giving us access to cell types that are otherwise difficult to obtain or follow through developmental time. Recently, systems genetics approaches have combined the power of genetic diversity with the differentiation potential of PSCs to identify genetic variants that influence gene expression regulation across different cellular contexts (Warren and Cowan 2018; Swanzey et al. 2021; Farbehi et al. 2024). For example, mouse embryonic stem cell lines (mESCs) derived from diverse strains have been used to study the maintenance of the pluripotent state, early lineage specification, and the stability of genomic imprinting (Skelly et al. 2020; Byers et al. 2022; Aydin et al. 2023; Parikh et al. 2024). Similarly, the influence of genetic variation on human induced PSCs (iPSCs) has been characterized at multiple molecular levels (Kilpinen et al. 2017; Panopoulos et al. 2017; Banovich et al. 2018; Mirauta et al. 2020) and throughout differentiation into various cell types (Cuomo et al. 2020; Jerber et al. 2021; Elorbany et al. 2022; Wells et al. 2023; Popp et al. 2024). Despite these examples, our understanding of the downstream consequences of pluripotent state variation on developmental trajectories and later stages of development remains largely undefined. To address this gap, we turned to a system where we could conduct systems genetics inquiry in both the pluripotent as well as a subsequent developmental stage using cells from the same donors.

PSCs can be differentiated into neural progenitor cells (NPCs) that can proliferate in culture continuously and have the capacity to further differentiate into specialized neuronal cell types such as neurons and astrocytes (Conti et al. 2005). Similar to their pluripotent ESC progenitors, NPCs represent a dynamic cell state maintained through external signaling molecules and growth factors in cell culture rather than a distinct cell type observed in vivo. NPCs are an increasingly important resource for cellular modeling of neurodevelopment and neurodegenerative disorders. For example, Young-Pearse and colleagues recently analyzed neuronal cell types differentiated from a set of 50 iPSC lines from the Religious Orders Study and Memory and Aging Project (ROSMAP) aging cohorts to identify a neuron-specific interaction between the Alzheimer’s Disease risk genes SORL1, APOE, and CLU (Lee et al. 2023). In a separate study, genetically diverse NPCs derived from human iPSCs were recently used to identify the genetic variants driving differences in susceptibility to Zika virus (Wells et al. 2023). Finally, another recent study profiled hPSC differentiation to mature neurons with single cell RNA sequencing (scRNA-seq) across a fine time course that captured progenitor-like cell states. The authors compared genetically identical lines in different cell states to identify molecular signatures that could predict differentiation efficiency into neuronal cells (Jerber et al. 2021). However, genetic diversity was limited in these human cell studies, and they lacked the sample sizes necessary to map trans eQTL interactions among genes that drive neural lineage specification.

Powerful mouse resources including the Diversity Outbred (DO) heterogeneous stock combine genetic variation from multiple parent strains in a randomized breeding design that minimizes population stratification and ensures balanced allele frequencies across the genome, providing optimal power for genetic mapping studies (Churchill et al. 2012). Recently, we applied a systems genetics approach to mESCs derived from DO mice to infer the complex gene regulatory network and genetic interactions that underlie ground state pluripotency (Skelly et al. 2020; Aydin et al. 2023). Specifically, we profiled chromatin accessibility, gene expression, and protein abundance in a large panel of DO mESCs and mapped thousands of quantitative trait loci (QTLs) that affected chromatin accessibility (caQTLs), transcript abundance (eQTLs) and protein abundance (pQTL) (Skelly et al. 2020; Aydin et al. 2023). In this study, we extend our systems genetics approach to characterize variation in transcript abundance and identify cell-state specific and conserved eQTLs in mouse NPCs derived from many of the same DO mESC lines used in our previous studies (Skelly et al. 2020; Aydin et al. 2023). In line with our findings in DO mESCs, we observe high expression variation among the DO mNPCs. Moreover, despite large expression differences overall between the mESC and mNPC cell states, we observed high co-variation in mESC and mNPC transcriptomes from the same DO donors. Genetic mapping identified thousands of significant local and distant eQTL, of which a quarter — mostly local eQTLs — were also detected previously in DO mESCs. Most distant eQTL, by contrast, were uniquely detected in mNPCs, including three eQTL hotspots on Chrs 1, 10, and 11. Targets of the Chr 1 hotspot were enriched for roles in chromosome segregation and the spindle assembly checkpoint (SAC). Cross cell-type mediation and partial correlation analysis identified two potential mediator genes of this hotspot, one expressed in the mNPCs (*Rnf152*) and the other earlier in the pluripotent mESCs (*Pign*). Both *Rnf152* and *Pign* have been implicated in cell-cycle regulation with *Pign* having a direct role in the SAC (Okamoto et al. 2020; Teye et al. 2021), suggesting that these genes drive variation in cell-cycle regulation in DO mNPCs. More broadly, our study demonstrates the power of applying a systems genetics approach across timepoints in a developmental trajectory, which allowed us to map eQTLs and link gene expression variation in one cell state to a causal variant affecting expression of a regulatory gene in a progenitor cell.

## Results

### Gene expression in genetically diverse mNPCs is highly variable and co-varies with gene expression in mESCs

To better understand how the abundant genetic variation segregating in the DO population affects gene expression during neural differentiation, we quantified the transcript abundance of 14,163 genes by RNA-seq in 186 DO mNPC lines (Figure 1A). We first performed principal component analysis (PCA) on mNPC gene expression (PCA-N) to identify the genes and pathways that are most variable across the DO mNPC lines. The first principal component (PC1-N, Figure S1A) explained 14% of expression variation among DO mNPCs, and functional enrichment analysis showed that PC1-N driver genes were overrepresented for those involved in the cell cycle, mRNA processing, translation, response to leukemia inhibitory factor (LIF), *in utero* embryonic development, and targets of transcription factors with roles in neural differentiation such as *Otx2* (Vernay et al. 2005) (Table S1). A closer look at the cell cycle-related processes revealed genes with roles in almost all of the cell cycle checkpoints, underscoring the proliferative nature of mNPCs under expansion culture conditions.

**Figure 1.**
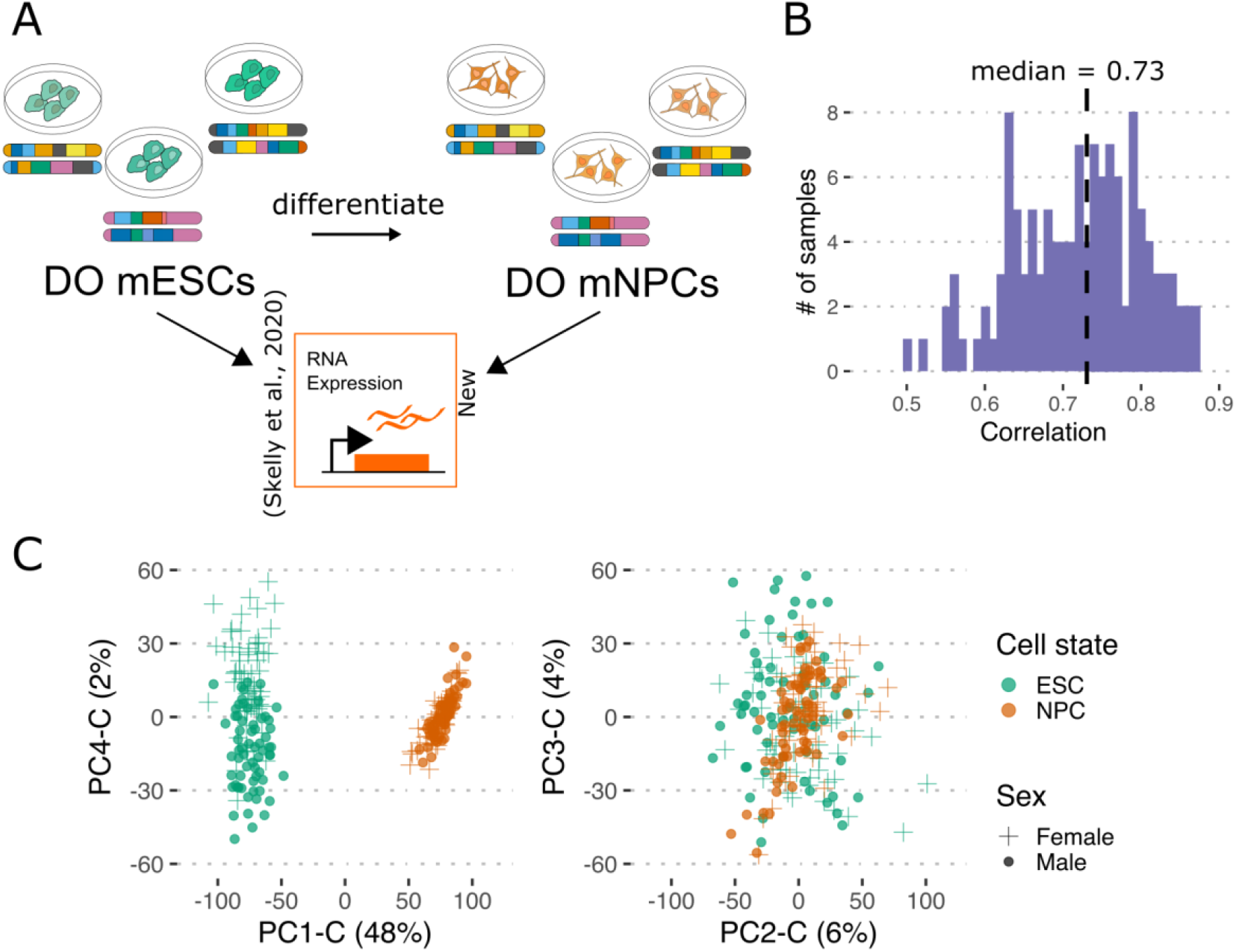
Variation in DO mNPC transcriptome and covariation to DO mESCs. (A) Experimental overview of the data. (B) Histogram of correlation coefficients between the ESC and NPC transcriptomes of 127 samples. (C) Principal component analysis results of the combined ESC, NPC transcriptomes of 127 samples and 12,095 genes.

Next, we sought to better understand how transcriptional variation in DO mNPC lines compared to variation in their progenitor pluripotent DO mESC lines. We previously characterized gene expression in mESCs from the same DO donor for 127 of the mNPC lines in the current study (Skelly et al. 2020). Over 12,000 genes were expressed in both cell states, and we observed significant covariation between the mESC and mNPC transcriptomes of these 127 cell lines (median intraline NPC-ESC correlation = 0.73, permutation test p < 2.2e-16; Figure 1B, Figure S1B), likely indicating a large role for genetic background in conferring expression variability. Similarly, we observed high concordance between the mean and variance of genes expressed in both cell states, with a correlation of 0.77 (p < 2.2e-16) and 0.74 (p < 2.2e-16), respectively (Figure S1C-D).

To discern the sources of expression variation that were unique to mNPCs or shared with mESCs, we performed PCA on the combined gene expression counts from both cell types (PCA-C). Cell state appeared to be the primary driver of expression variation in the combined dataset (PC1-C, 48%; Figure 1C), despite the high within-line co-variation of expression in mNPCs and mESCs.

Functional enrichment analysis of PC1-C implicated ubiquitin mediated protein degradation, nervous system development, signaling pathways regulating pluripotency, and cellular differentiation as the major drivers of gene expression variation between the mESC and mNPC lines (Table S2). Of note, PC1-C showed high agreement with PC1-N values (cor = -0.9) for the 127 DO mNPC lines included in both analyses. However, we find little overlap in the genes contributing to PC1-N and PC1-C, with only 39 genes shared among 709 driver genes for PC1-N and 605 driver genes for PC1-C. While PC1-C clearly corresponded to cell state expression differences, no other PC in the combined dataset appeared to distinguish between cell states; indeed, PC2-C and PC3-C did not show any separation, exhibiting a range of values that spanned both cell states, and likely capturing variation common to both (Figure 1C). Drivers of PC2-C and PC3-C were enriched for genes involved in essential cellular processes including the mitotic cell cycle, mitochondrial cell function, mRNA processing, membrane lipid metabolism and chromatin organization (Table S2). PC4-C appears to differentiate mESC lines by their chromosomal sex, which was previously observed and attributed to the two active X chromosomes in the pluripotent mESC state (Skelly et al. 2020; Aydin et al. 2023). This strong sex effect is not present in the mNPCs where the second X chromosome is known to be inactivated (Figure 1C). Variance decomposition analysis confirms this difference in sex effect on gene expression in mESCs and mNPCs (Figure S1E).

To better characterize biological processes that were differentially regulated across the cell states, we performed differential gene expression analysis. We identified 1,409 upregulated and 773 downregulated genes in mNPCs relative to mESCs (adjusted p < 0.05, abs(log2 fold change) > 2). Genes upregulated in mNPCs were enriched for biological processes including nervous system development and pathways involved in neural differentiation such as WNT signaling. Conversely, genes downregulated in mNPCs were enriched for biological processes including ribosome biogenesis and pathways regulating pluripotency such as response to LIF (Table S3), confirming the PCA results.

### Genetic architecture of the mNPC transcriptome

To further characterize the sources of interline expression differences in mNPCs, we estimated narrow sense heritability. We estimated the transcript abundance of over 90% of expressed genes in mNPCs to be heritable (median *h*^*2*^ = 0.23) — confirming that genetic diversity among DO mNPCs is a major driver of transcriptome variation. Genetic mapping identified 4,060 significant eQTL (LOD > 7.5, alpha = 0.05, FDR = 0.075), with almost one-third of expressed transcripts having one or more significant eQTL (Table S4, Figure 2A). The majority of eQTL are local (n = 2,998), meaning the genetic variant impacting the abundance of the transcript is within close proximity (± 10Mb) to the gene itself. The remaining quarter are distant eQTL (n = 1,062), where the peak is further away from the gene or on a different chromosome and likely influences the target gene’s expression indirectly through direct effects on the expression or function of another “mediator” gene. Similar to previous eQTL analyses, we observed that many distant eQTL colocalized to eQTL “hotspots”. We detected three such eQTL hotspots on Chrs 1, 10, and 11 that were unique to mNPCs (Figure 2B, Table S5).

**Figure 2.**
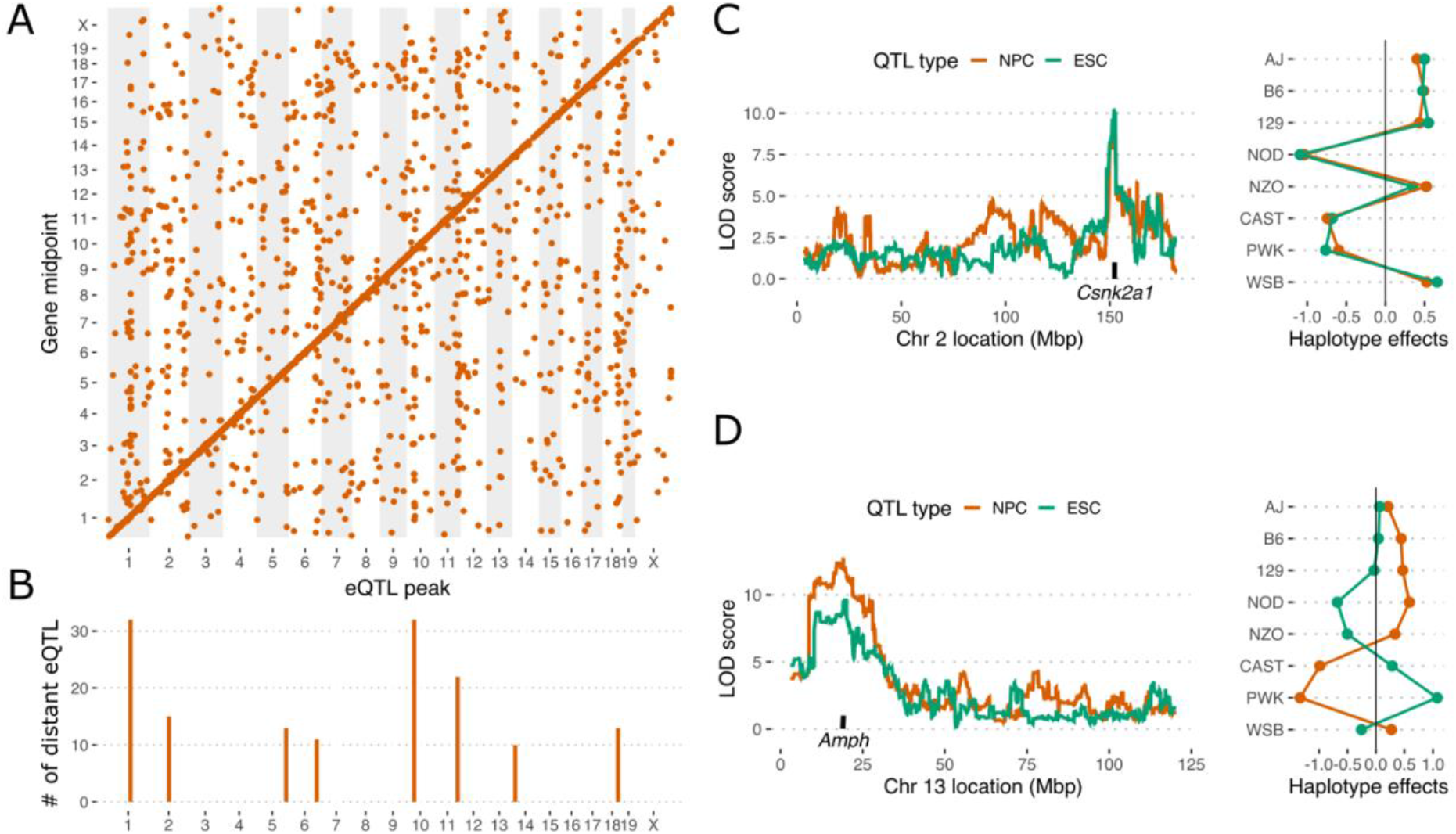
Genetic architecture of DO mNPC transcriptomes. (A) Map of DO mNPC eQTL containing 4,060 significant peaks (LOD > 7.5) from 3,806 unique genes. (B) eQTL hotspots identified across the genome. (C) eQTL scan for *Csnk2a1* gene on the left and the haplotype effects at the eQTL peak on Chr 2 on the right. (D) eQTL scan for *Amph* gene on the left and the haplotype effects at the eQTL peak on Chr 13 on the right.

Interestingly, we also mapped a significant QTL for PC1-N values (LOD > 7.11, alpha = 0.1) to the same hotspot on Chr 1, suggesting that this hotspot is a causal driver of global transcriptional variation in mNPCs (Figure S2A).

Next, we compared our mNPC eQTL map to the mESC eQTL map from our previous study (Skelly et al. 2020). Over a quarter of eQTL in mNPCs are also detected in mESCs and show concordant allele effects (abs(cor) > 0.7 and adjusted p < 0.1; n = 1,113 concordant eQTL / 4,060 total eQTL, Figure S2A-B); these concordant eQTL are primarily local variants (963 local eQTL out of 1,113 total concordant eQTL). For example, *Csnk2a1* has a shared local eQTL with highly similar effects in both mNPCs and mESCs (Figure 2C). *Csnk2a1* is known to regulate various cellular processes including WNT signaling, cell cycle progression, apoptosis and transcription, and mutations in this gene have recently been linked to neurodevelopmental abnormalities in humans (Okur et al. 2016). Of note, we also map concordant local eQTL with opposing allele effects on gene expression in mNPCs and mESCs. Although much fewer in number, one such example is *Amph* (Figure 2D), a gene involved in synaptic vesicle endocytosis where loss of expression leads to cognitive deficits in mouse models (Paolo et al. 2002; Zhang et al. 2021). The remaining three quarters of eQTL are unique to mNPCs (n = 2,947 unique / 4,060 total eQTL) and affect target genes that are enriched for roles in cell-cell signaling and nervous system development (Table S6).

### Cross-cell type mediation analysis uncovers the influence of genetic variation across cell states

To characterize the eQTL hotspots uniquely observed in DO mNPCs, we performed functional enrichment analysis to identify affected pathways and mediation analysis to identify candidate regulatory genes in the hotspot regions. For these analyses, we relaxed our inclusion criteria to include all target genes that mapped with a suggestive eQTL (LOD > 6) in the three hotspot regions on Chrs 1, 10, and 11. Targets of the Chr1 hotspot were significantly enriched for genes involved in mRNA processing, DNA repair, chromatin organization, protein degradation and cell cycle (adjusted p-value< 0.01; Table S7). Genes involved in protein processing in the ER were significantly overrepresented for the hotspot on Chr 10, while the Chr 11 hotspot did not show any significant functional overrepresentation. Mediation analysis failed to identify strong candidate genes underlying the Chrs 10 and 11 eQTL, suggesting that these hotspots may stem from genetic variant(s) affecting abundance of an RNA not profiled (e.g., miRNA), a post-transcriptional modification of RNA, function or stability of a protein, or any mechanism that does affect the transcript abundance of the mediator gene. We excluded these loci from further analysis.

Measurement error — including read mapping errors in RNA-seq datasets — can adversely affect individual gene expression estimates and introduce noise into eQTL mapping. Dimensionality reduction techniques like PCA are an effective way to summarize the main component of expression variation among eQTL hotspot target genes to infer founder allele effects and identify candidate mediator genes. Accordingly, we performed PCA on the transcript abundance of hotspot target genes and mapped QTL for the eigengene value (Eigen-Chr1). As expected, Eigen-Chr1 mapped with a significant QTL to the Chr 1 hotspot and exhibited founder haplotype effects similar to the observed individual target gene eQTL. Specifically, DO mNPC lines that inherited the Chr 1 hotspot region from B6, 129, NOD, or CAST founder strains showed similar expression of the eQTL target genes, while those lines that inherited this locus from AJ, NZO, PWK, or WSB showed a different pattern of gene expression (Figure 3A, B). Of note, the PC1-N Chr 1 QTL also showed the same 4:4 split in founder haplotype effects (Figure S3A). PC1-N values showed high agreement to Eigen-Chr1 (cor = -0.99), which likely reflects the significant overlap between the individual gene targets of the Chr 1 hotspot (overlap n = 193 / 322) and the gene drivers of PC1-N (overlap n = 193 / 709, hypergeometric p = 1e-86).

**Figure 3.**
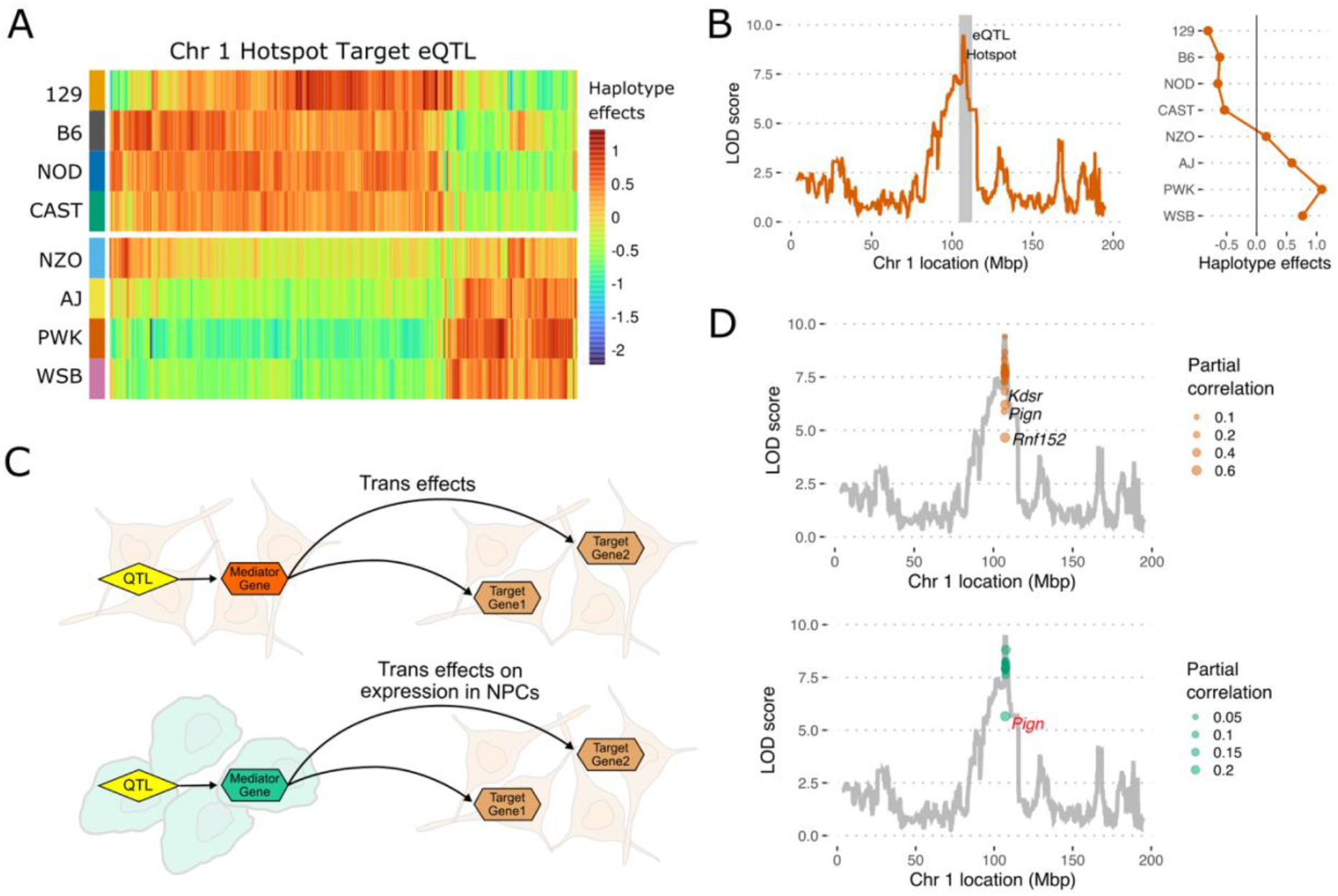
Details of the Chr 1 eQTL hotspot. (A) Heatmap of founder haplotype effects at the eQTL peaks of the target eQTL (LOD >6, n = 322) within the Chr 1 hotspot. (B) QTL scan of PC1-T of the Chr 1 target eQTL on the left, founder haplotype effects at the PC1-T QTL peak on the right. (C) Cartoon depicting mediation analysis using ESC and NPC expression to identify potential regulators of an eQTL hotspot. (D) In gray, PC1-T QTL scan is plotted overlaid with the new LOD scores obtained after mediation with genes in the region. Red gene name depicts genome wide significance for the mediation LOD drop.

Given the large physical distance between eQTL hotspots and their target genes, it is generally assumed that the trans effects of these variants are conferred via direct proximal effects (i.e. cis effects) on the expression or function of a regulatory gene in the hotspot region. In cases where the causal genetic variant alters the expression level of the regulatory gene, mediation analysis can be a powerful tool to identify this “mediator” gene (Chick et al. 2016; Skelly et al. 2020). Importantly, an eQTL hotspot in one cell state (e.g. mNPCs) could stem from a genetic variant(s) that influences the expression of a regulatory gene in that same cell state *or* in a progenitor stem cell (e.g. mESCs). We performed mediation analysis on the Eigen-Chr1 values to identify potential mediators of the Chr 1 hotspot target genes, searching first in the mNPC expression data and then in the mESC expression data (Figure 3C). Initially, we used a standard regression model to identify this potential local regulator within the Chr 1 hotspot (Chick et al. 2016). To gain more insight into the relationship between candidate mediators and the Chr 1 hotspot, we also performed partial correlation analysis — assessing the correlation of expression levels between candidate mediators and target genes after accounting for the observed effect of the distant eQTL — with the expectation that a true causal mediator should still co-vary in expression with the target genes even in the absence of the observed QTL effect. We calculated the partial correlation between Eigen-Chr1 and the expression of candidate mediators after accounting for the genetic ancestry at the QTL peak.

Mediation analysis with the mNPC expression data identified *Rnf152* as the best candidate mNPC mediator in the Chr 1 hotspot. That is, regressing out *Rnf152* expression in mNPCs caused the largest LOD drop in the most target eQTL and the Eigen-Chr1 QTL (Figure 3D, Figure S3B). None of the candidate mediators, including *Rnf152*, were statistically significant as assessed by genome-wide scaling, but *Rnf152* showed the highest partial correlation to Eigen-Chr1 after regressing out the QTL effect (correlation = 0.6, adjusted p = 3.5e-12). *Rnf152* encodes a ubiquitin ligase residing in the lysosome; it is reported to be involved in mammalian target of rapamycin (mTOR) signaling and has been shown to regulate autophagy and cell proliferation (Deng et al. 2015; Okamoto et al. 2020). *Rnf152* has a local mNPC eQTL with a split in founder allele effects similar to the eQTL hotspot, consistent with its role as a Chr 1 hotspot mediator. (Figure S3C).

Next, we expanded our mediation analysis across cell states to identify potential mediators of the mNPC Chr 1 hotspot in the mESC expression data. We identified *Pign* as the best candidate mESC mediator for the mNPC Chr 1 hotspot, causing the largest LOD drops in the most target eQTL and the Eigen-Chr1 QTL. Further, mESC *Pign* expression showed the highest partial correlation to Eigen-Chr1 after regressing out the Chr 1 eQTL effect (correlation = 0.3, adjusted p = 0.1) (Figure 3D, Figure S3B). The best mNPC mediator, *Rnf152*, was found to be expressed at very low levels in mESCs and did not pass our filtering threshold — consistent with previous reports (Lienert et al. 2011). *Pign* encodes for an enzyme involved in glycosylphosphatidylinositol (GPI)-anchor biosynthesis, and more recently was linked to cell cycle regulation through interaction with the spindle assembly checkpoint (SAC) complex (Teye et al. 2021). In DO mESCs and mNPCs, *Pign* has a significant local eQTL with effects that match the 4:4 founder allele split observed for the eQTL hotspot (Figure S3D). Importantly, target genes of the Chr 1 hotspot include those involved in proper chromosome segregation at mitosis such as *Mad2l1* (MAD2), a member of the SAC complex (Dobles et al. 2000).

## Discussion

Over the past two decades, genome-wide association studies (GWAS) in large human cohorts have identified thousands of common variants associated with variation in numerous complex traits and disease phenotypes (Abdellaoui et al. 2023). Many of these variants likely influence adult phenotypes through direct effects on gene expression or cell differentiation early in development, making them difficult to assay directly in humans (Maurano et al. 2012). In this study, we focused on neural differentiation, and sought to better understand how genetic variation can influence gene expression using a large panel of mouse neural progenitor cells derived from the powerful Diversity Outbred heterogeneous stock. We analyzed interline variation in mNPC gene expression, performed eQTL mapping and mediation analysis, and compared our results to pluripotent mESC lines from the same donors. We identified genetic variation to be a significant driver of gene expression variability in DO mNPCs. Although they represent very different cellular states and this difference is clearly evident in their transcriptomes, genetically identical mESC and mNPC lines showed high covariance in gene expression (median intraline correlation = 0.73) — indeed, considerably higher agreement than we observed between transcript and protein abundance within genetically identical mESC lines (median intraline correlation = 0.36) (Aydin et al. 2023). Gene expression differences between cell states highlight known differentiation pathways, as genes involved in nervous system development and differentiation are upregulated in mNPCs while genes with roles in pluripotency maintenance are downregulated in mNPCs relative to mESCs.

Meanwhile, genes involved in essential cellular functions like the cell cycle, mitochondrial function, RNA processing, and membrane lipid metabolism were found to be variably expressed in both cell states. Given that mESCs and mNPCs are both self-renewing in culture, and given the dynamics and requirements of actively dividing cells, we were not surprised to observe high expression variability for genes involved in cell cycle regulation and cellular metabolism.

Expression QTL mapping added context to these expression patterns and uncovered thousands of genetic variants responsible for their expression variation. We found that over a quarter of significant eQTL detected in mNPCs were also mapped in mESCs and showed similar allele effects.

These concordant eQTL were primarily locally-acting and affected many genes with essential cellular functions. Population variation at these loci likely accounts for much of the high expression covariation observed in mNPCs and mESCs from the same DO donor. The remaining three quarters of mNPC eQTL appear unique to this cell state; of note, a similar proportion of mESC eQTL were also cell state-specific (4,854 / 6,069 or 80%). Many of these cell state-specific distant eQTL co-localize to genomic hotspots and likely play significant roles in cell lineage specification. For example, the previously identified Chr 15 eQTL hotspot regulating pluripotency maintenance is only observed in DO mESCs (Skelly et al. 2020; Aydin et al. 2023), while the hotspots on Chrs 1, 10, and 11 are only observed in DO mNPCs. Further, we found that genes involved in cell-cell signaling and nervous system development are significantly overrepresented among the unique distant eQTL in DO mNPCs. Together, these results highlight the abundance and widespread distribution of expression-modulating variants in genetically diverse populations, underscore a role for conserved eQTL in driving interindividual variation within cell states, and provide further support for the importance of distant eQTL to lineage specification driving cell state-specific functions.

We identified three eQTL hotspots on Chrs 1, 10, and 11 with distinct genetic effects in DO mNPCs. Furthermore, mediation analysis helped us identify potential regulators for the hotspot on Chr 1. One of the top candidates, *Rnf152*, is a ubiquitin ligase that regulates autophagy. Although the drop in LOD scores obtained through mediation analysis did not achieve genome-wide significance for the target eQTL, *Rnf152* showed the highest partial correlation to the PC1 of the expression of target genes when controlled for the founder strain ancestry at the QTL peak. Interestingly, the other candidate regulator, *Pign*, was identified from the ESC state. *Pign* was shown to regulate the spindle assembly checkpoint (SAC) by forming a complex with the SAC proteins MAD1, MAD2, and the mitotic kinase MPS1 where loss of *Pign* expression led to an increase in errors in chromosome segregation and dysregulation of the SAC (Teye et al. 2021). In addition, *Pign* is linked to a variety of human disorders including some with neurological symptoms like developmental delay, epilepsy and seizures (Maydan et al. 2011). Although neither *Rnf152* nor *Pign* are transcription factors and are unlikely to directly regulate gene expression, they could be indirectly regulating the expression of downstream target genes through cellular signaling and influencing the progression of the cell cycle. *Rnf152* and *Pign* are separated by less than 200kb on Chr 1, and we identified four candidate SNPs in the region that match the observed 4:4 strain allele effect (rs31971829, rs47676250, rs37516825, rs47457121) and may influence expression of one or both genes. Future experiments will seek to validate the causal variant(s) underlying this mNPC hotspot eQTL and establish the cell state of origin (i.e., mNPC or mESC) for its direct effects.

In this study, we obtain the first comprehensive look at the variation in the transcriptomes of a genetically diverse set of mNPCs. In addition, we characterized the influence of genetic variation across cell states using a large panel of genetically identical mESC and mNPC lines. Our experimental design allowed us to trace the impacts of genetic variation across two states in a developmental trajectory, which showed that primarily local genetic effects are shared between cell states. However, trans effects, even ones with large influences (e.g. *Lifr* allele in mESCs (Skelly et al. 2020)), are not conserved. Combined with the overrepresentation of neural development genes among those with unique distant eQTL in mNPCs, our results emphasize the specialized regulation of gene expression across cell states.

Finally, we acknowledge a number of study limitations that temper the strength of our conclusions. Although comparable in size to other published eQTL studies, our DO mNPC panel is not optimally powered to detect variants that confer only subtle effects. Given that essential regulatory genes may not tolerate large effect variants, we concede that our inability to detect small effect eQTL may cause us to miss important genetic interactions. Mediation analysis is a powerful tool but limited to cases where the causal variant(s) underlying the distant eQTL modulates the transcript abundance of the mediator gene. Distant eQTL that stem from causal variants that influence post-transcriptional regulation, protein function or stability, or any other mechanism that does not impact transcript abundance of the mediator gene will be missed by mediation analysis. Similarly, partial correlation between genes does not necessarily indicate a direct regulatory or functional interaction.

## Materials and Methods

### Neural differentiation of DO mESCs

Diversity Outbred ES cell lines were produced as previously described (Skelly et al. 2020). Neural stem cell lines were produced by differentiation of ES cell lines following previously established protocols (Pollard et al. 2006). Briefly, cryopreserved ES lines were thawed and carried for 3 to 6 passages prior to differentiation. ES cells were trypsinized and 50K to 75K cells were plated per well of a laminin treated 12 well plate in NS Media with FGF2 and EGF (Pollard et al. 2006). The medium was changed daily for 8 days, then detached with accutase, washed with NS media and replated into laminin coated 6 well plates with 2mls per well of NS media. Neural stem cells were expanded in clusters for 3 days then serially expanded into 6cm and then 10cm dishes until subconfluent. NS cells were passaged for an additional 3 weeks and then cryopreserved. NS lines were screened positive for neural stem cell markers such as nestin and GLAST, and negative for ES cell markers such as SSEA1. The lines were further validated by differentiation to terminal neural cell types such as astrocytes and neurons, as demonstrated by flow cytometry after cell type specific staining.

### Diversity Outbred mNPC RNA-seq

Total RNA was isolated from each of 186 DO mNPC lines and quantitated by paired-end RNA sequencing. Briefly, for each mNPC line, one 15cm dish of cells was grown to near confluence, washed 3x with PBS, and mechanically harvested to yield 10M cells. About 100k cells from each frozen cell pellet were used for RNA sequencing. Next, total RNA was extracted using the Quick-RNA 96 well format kit (Zymo Research) with in-column DNase treatment. Sequencing libraries were prepared by Akesogen using the TruSeq Stranded mRNA HT kit (Illumina, Cat no. 20020595) and included ribosomal RNA reduction and poly-A selection, enzymatic fragmentation, cDNA synthesis from random hexamer priming, adaptor ligation and PCR amplification steps to generate indexed, stranded mRNA-seq libraries. Libraries were checked for quality and quantitated with the Agilent Bioanalyzer, and samples that failed QC were repeated starting from the cryovial stage.

Finally, pooled libraries were sequenced on the NextSeq platform (Illumina) using the NextSeq 500/550 High Output v2 150-cycle kits (Illumina, Cat no. FC-404-2002). To minimize technical variation, samples were randomly assigned to lanes prior to sample processing steps, barcoded, and multiplexed at 16 samples per flow cell, yielding 6M-55M 2×75bp paired-end (PE) reads per sample.

We aligned paired-end 75bp reads with bowtie v1.1.2 (Langmead et al. 2009) to a pooled “8-way” transcriptome containing strain-specific isoform sequences from all eight DO founder strains as previously described (Skelly et al. 2020). In order to identify and correct sample mix-ups, we inferred sample genotypes from the RNA-seq data using GBRS v0.1.6 (Choi et al. 2020) and compared GBRS-derived genotypes to our DNA genotypes obtained from GigaMUGA arrays. Correlations between genotypes inferred from GBRS and GigaMUGA for the same sample were typically on the order of 0.8-0.9 whereas for different samples they were less than 0.5. We resolved 17 sample mix ups for DO mNPC lines showing an incongruent genotype. For resolving multi-mapping reads and quantifying transcript- and gene-level expression counts, we utilized EMASE as implemented in GBRS v0.1.6 (Raghupathy et al. 2018; Choi et al. 2020). We filtered genes with a median TPM (transcripts per million) value smaller than 0.5 or zero value (i.e., not expressed) in more than half of the samples. To account for differences in library size, we normalized gene-level counts to the upper quartile value then applied the ComBAT function from R/sva package to remove batch effects caused by library preparation (Johnson et al. 2007). We transformed normalized and batch corrected values to rank normal scores using rankZ normalization (Gatti et al. 2014). Gene annotations such as MGI symbol, gene location and gene biotype were added using v84 of the Ensembl database.

### Diversity Outbred mESC RNA-seq

Raw RNA-seq data was retrieved from ArrayExpress (E-MTAB-7728) and analyzed same as to the DO mNPC RNA-Seq data.

### Statistical analysis

All analyses and figures were generated with the R statistical programming language (R Core Team 2023). Unless otherwise stated, R/tidyr package was used for data processing, R/ggplot2 for plotting, and R/pheatmap for heatmap plots.

### Functional enrichment analysis

We performed functional enrichment analysis using the ‘gost’ function in the gProfiler2 package (Raudvere et al. 2019; Kolberg et al. 2020) by controlling the version using ‘set_base_url(set_base_url(“https://biit.cs.ut.ee/gprofiler_archive3/e102_eg49_p15/“))’ in R using an appropriate universal background on a case-by-case basis and ‘fdr’ option for p-value correction. For example, we used the list of all genes that are expressed in NPCs for enrichment analysis of the DO mNPC Chr 1 hotspot target eQTL. For Gene Set Enrichment Analysis (GSEA) we utilized the R/fgsea package (Korotkevich et al. 2021) with gene sets that belong to the Gene Ontology Biological Processes (GO:BP) subcategory in the molecular signatures database (MSigDB) and 10,000 permutations.

### Differential expression analysis

Differential expression was calculated for the 12,095 genes expressed in both mESCs and mNPCs from the same DO donor (n = 127 lines total). In addition to the removal of the batch effects, sex effects were also removed before the comparison in both data sets using ComBAT as implemented in the R/sva package (Johnson et al. 2007). We compared gene expression using the Wilcoxon Rank Sum test, implemented as the ‘wilcox.test’ function in the R/rstatix package, and adjusted raw p-values to correct for multiple testing using the Benjamini-Hochberg method “BH”. The log2 fold change was calculated by taking the log2 of the ratio between the mean expression of each gene in mNPCs over mESCs.

### Quantitative Trait Locus mapping

Genetic mapping was performed using the R/qtl2 package (Broman et al. 2019) with wrapper functions utilizing parallelization for efficient large-scale eQTL analysis in R/QTLretrievR (https://github.com/deweyhannah/QTLretrievR). Briefly, eQTL were mapped with a linear mixed model — implemented as the ‘scan1’ function in R/qtl2 — using the upper quartile normalized, batch corrected, rankZ transformed gene expression values, with chromosomal sex included as an additive covariate and the Leave One Chromosome Out (loco) option used for kinship correction(Yang et al. 2014). To estimate genome-wide significance, we permuted genotypes 1000 times while maintaining the relationship between the phenotype and covariates. For each permutation we retained the maximum LOD score to generate a null distribution for the test statistic (Churchill and Doerge 1994). To calculate thresholds for eQTL, we repeated this permutation strategy for all transcripts and estimated a significance cutoff at LOD > 7.5 (alpha = 0.05), and a suggestive cutoff at LOD > 6. False discovery rates (q-values) were determined for each permutation-derived p-value with R/qvalue software, using the bootstrap method to estimate pi0 and the default lambda tuning parameters (Storey et al. 2004) and our LOD threshold of 7.5 corresponds to an FDR of 0.075. We call a QTL ‘local’ if the QTL peak is within ±10 Mbp to the midpoint of its corresponding gene and ‘distal’ if otherwise.

Founder allele effects were estimated as best linear unbiased predictors (BLUPs) at the QTL peak using ‘scan1blup’ function in R/qtl2 package. To identify overlaps with significant NPC eQTL, we used a relaxed threshold of LOD > 5 for ESC eQTL. They were classified as shared if the eQTL peaks were within +/-5 Mb of each other and the correlation between haplotype effects was significant (adjusted p-value < 0.1).

For hotspot calling, we first identified distal eQTL that reach genome-wide permutation-based threshold (p < 0.05; LOD 7.5). Next, we applied a sliding window method to identify hotspots as described in (Skelly et al. 2020). Briefly, we counted the number of distal eQTL within 1cM windows (0.25 cM shift) across the genome and selected the top 0.5% of bins with the most distant eQTL (0.5% bin threshold 9 distant eQTLs). Final coordinates for each hotspot were determined using the Bioconductor package ‘GenomicRanges’ to merge adjacent bins into a single region (Lawrence et al. 2013). Each hotspot contains >20 significant distant eQTL (>150 suggestive, LOD > 6) and shows distinct allelic effect patterns.

### Partial correlation analysis

We used the ‘pcor’ function from the R/ppcor package (Kim 2015) to calculate the partial correlation between the expression of the mediator gene and the first principal component of the expression of genes with the target eQTL (Eigen-Chr1) on Chr 1. To control for the genetic effects in the partial correlation analysis, we classified the genotypes at the Eigen-Chr1 QTL peak based on their ancestry from the 8 DO founder strains. Given the 4:4 split at the Eigen-Chr1 QTL peak, if the cell line showed homozygous ancestry from any of the strains in the B6/129/NOC/CAST group they were classified as “Ref”, homozygous ancestry from any of the AJ/NZO/PWK/WSB strain group they were classified as “Alt”, and if they showed ancestry from both groups were classified as “Het”. We calculated the partial correlation for all the candidate mediators within ± 10Mb of the Eigen-Chr1 QTL peak using both gene expression from DO mESCs and mNPCs.

### Mediation analysis

We performed mediation analysis to identify candidate causal genes in mNPC eQTL hotspots. Mediation analysis was performed using the R/intermediate package (https://github.com/simecek/intermediate) by regressing target eQTL on the expression of a candidate mediator in the QTL and adjusting for covariates. We applied the ‘double-lod-diff’ method to reduce the effects of missing values. For mediation of QTL with the matching cell state we used the full sample set, e.g., eQTL mediation by NPC transcripts were done using all the 186 samples. Mediation across cell states was performed with mESC expression data from the common set of 127 DO donors. To assess the significance of a LOD drop, we mediated the QTL against all expressed genes genome-wide in that cell state to establish a null distribution of LOD score drops, converted the recorded LOD scores to normal scores, and checked if the score fell below 4 standard deviations from the mean of the null distribution. Mediators were further filtered to narrow down top candidates to include genes with midpoints that are found within ± 10Mb of the QTL peak.

## Data availability

The raw and processed RNAseq files are deposited to the Sequencing Read Archive and Gene Expression Omnibus (GSE285231).

## Acknowledgments

We would like to thank all members of the Munger laboratory for comments and discussion. We would also like to thank Ann Wells, Greg Carter, and other members of the Carter laboratory for their helpful feedback on the study and manuscript, and Charles Farber and Emily Farber of The University of Virginia Genome Analysis and Technology Core for providing sequencing services and support. This work was supported by the National Institutes of Health (NIH) grants GM133495 to S.C.M.; GM133724 to C.L.B.; HHSN273201500196P to T.C.; R24OD030037 to L.G.R, C.L.B., and S.C.M.

## Author Notes

## Conflict of Interest

T.C. has an equity interest in Predictive Biology, Inc. All other authors report no conflicts of interest.

## Supplemental Figures

**Figure S1.**
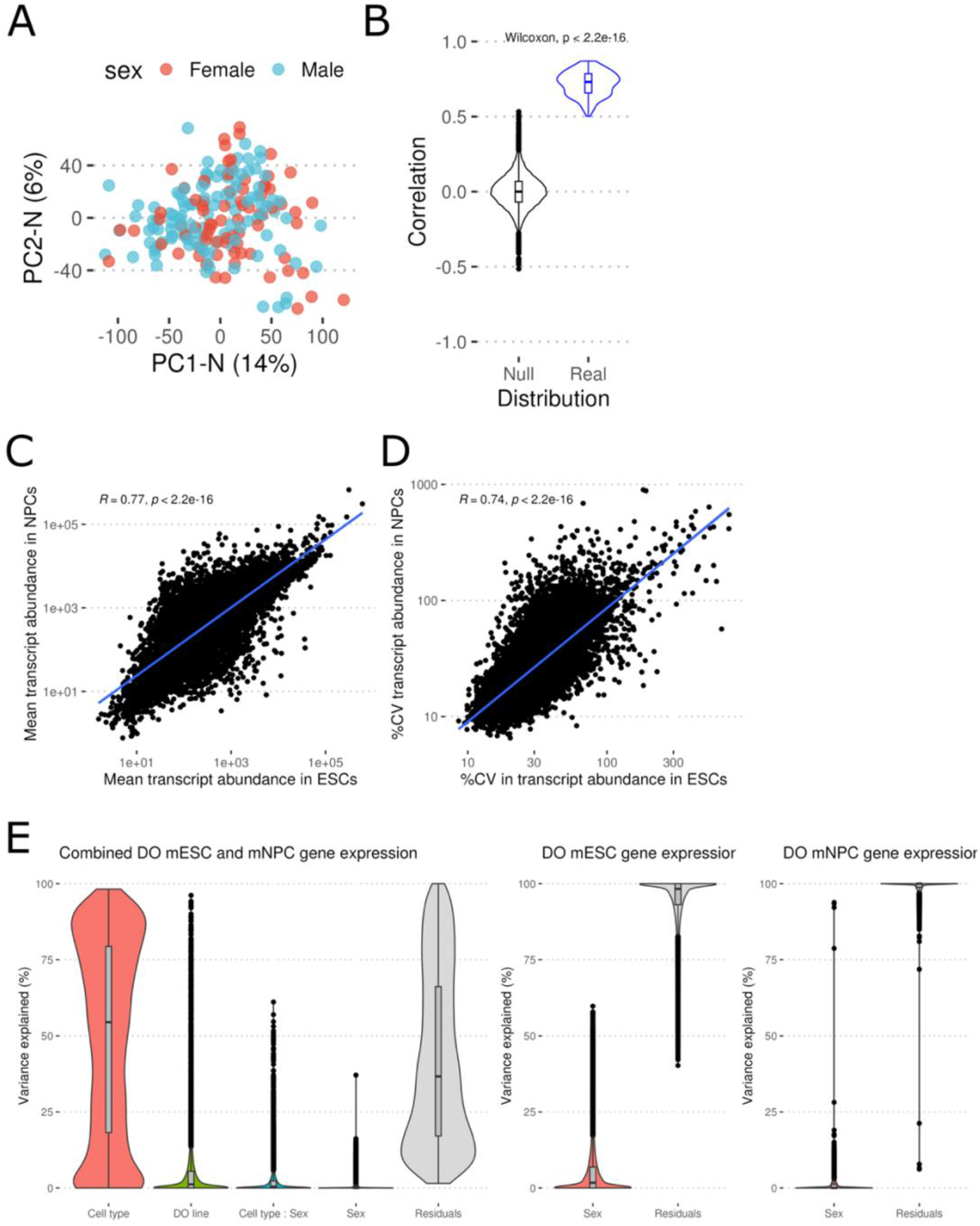
(A) Principal component analysis results of the DO mNPC data. (B) Genetically identical cell lines show significantly higher correlation than what is expected by chance between the ESC and NPC transcriptomes. Violin plots overlaid with boxplots depicting the distribution of Pearson correlation coefficients between the transcriptomes of 127 genetically identical mESC and mNPCs (blue) and the null distribution generated through 1000 permutations where the sample names are randomized (black). (C-D) Scatterplots showing mean and coefficient of variation (% CV) for transcript abundance for genes with measurements in both mESCs and mNPCs. (E) Violin plots overlaid with boxplots showing the variance partition analysis results in the combined and individual DO mESC, mNPC transcriptomes.

**Figure S2.**
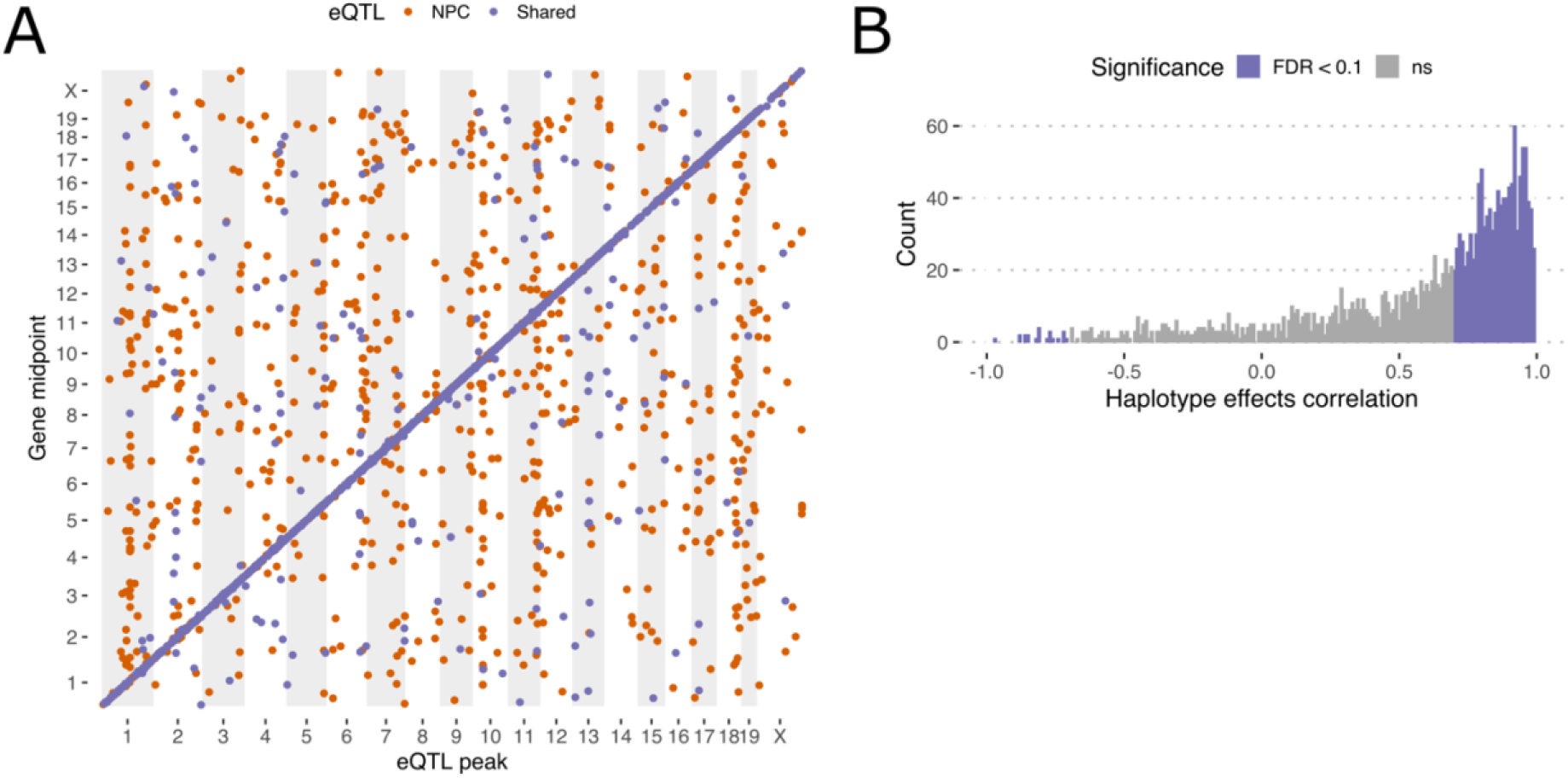
(A) Genetic mapping identifies 2,143 eQTL unique to NPCs and 2,529 eQTL shared between ESC and NPCs. The location of the eQTL is plotted on the x axis against the midpoint of the gene on the y axis. (B) Most co-mapping eQTL show high agreement in their haplotype effects. Histogram of pairwise correlation coefficients between inferred allele effects from eQTL from ESC and NPC scans for all genes with co-mapping QTLs. Bars are colored by significance of the correlation.

**Figure S3.**
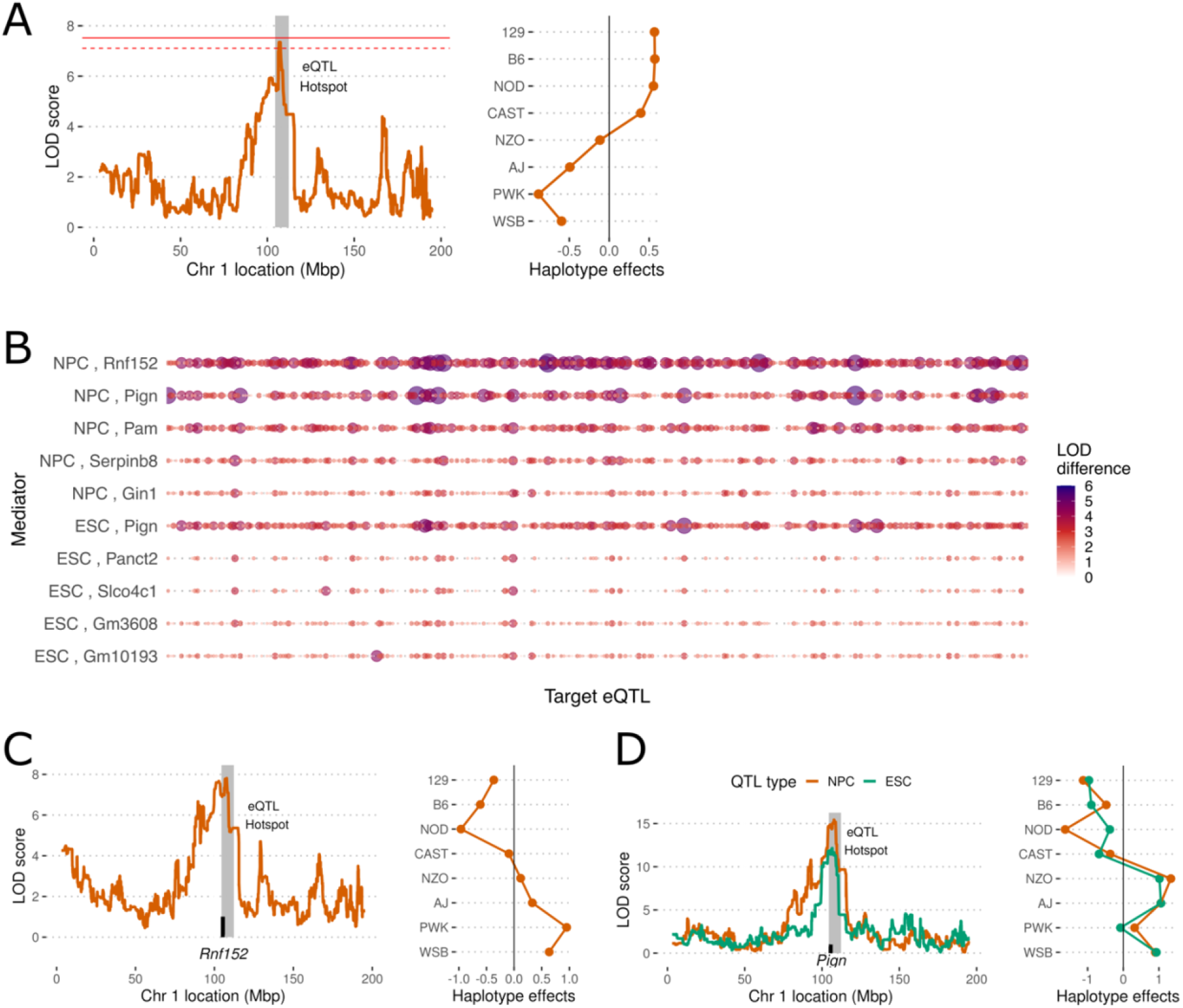
(A) Genetic mapping with PC1-N identifies a significant QTL on chromosome 1 with similar haplotype effects to the eQTL hotspot on chromosome 1. The red lines correspond to LOD thresholds for alpha = 0.05 (solid) and alpha = 0.1 (dashed). On the right, inferred haplotype effects at the QTL peak is plotted. (B) drops obtained from mediation analysis using NPC and ESC expression is plotted for the chromosome 1 target eQTL, showing results for the 5 best mediator genes within 10Mb of the eQTL hotspot. (C) Genome scan of *Rnf152* expression in NPCs show a significant local eQTL peak on chromosome 1 with inferred haplotype effects at the peak plotted on the right. (D) Genome scan of *Pign* expression in ESCs (green) and NPCs (orange) show a significant local eQTL peak on chromosome 1. The inferred haplotype effects at the peak are plotted on the right.

## Supplemental Tables

**Table S1**. List of biological pathways and processes over-represented in DO mNPCs PCA PC1-N.

**Table S2**. List of biological pathways and processes over-represented in PC1-C, PC2-C and PC3-C drivers from the combined data PCA.

**Table S3**. List of biological pathways and processes enriched in genes differentially expressed between DO mNPCs and mESCs.

**Table S4**. List of all significant DO mNPC eQTL.

**Table S5**. Details of DO mNPC eQTL hotspots and target genes.

**Table S6**. List of biological pathways and processes over-represented in shared and unique DO mNPC eQTL.

**Table S7**. Biological pathways and processes over-represented in target genes of chromosome 1 and 10 eQTL hotspots.

